# *PIK3R3* is a candidate regulator of platelet count in people of Bangladeshi ancestry

**DOI:** 10.1101/2022.11.11.516129

**Authors:** Kate Burley, Lucy Fitzgibbon, David van Heel, Genes & Health Research Team, Dragana Vuckovic, Andrew D Mumford

## Abstract

Platelets are mediators of cardiovascular disease and are regulated by complex sets of interacting genes. To reveal new regulatory loci for platelet count (PLT), we performed genome-wide association studies (GWAS) in 20,218 Bangladeshi (BAN) and 9,198 Pakistani (PAK) individuals from the Genes and Health study. Most significantly associated loci (p <5 ×10^−8^) replicated findings in prior transethnic GWAS. However, the BAN locus defined by rs946528 (chr1:46019890) did not associate with PLT in the PAK analysis but was in the same linkage disequilibrium block as PLT-associated variants in prior East Asian GWAS. The single independent association signal was refined to a 95% credible set of 343 variants spanning eight coding genes. Functional annotation, mapping to megakaryocyte regulatory regions and colocalization with whole blood eQTLs identified the most likely mediator of the PLT phenotype to be *PIK3R3* encoding a regulator of phosphoinositol 3-kinase, widely linked elsewhere to adverse cardiovascular phenotypes.

## Introduction

Blood platelets are essential mediators of haemostasis and are implicated in multiple cardiovascular and inflammatory disorders. Circulating platelet count (PLT) is an independent predictor of morbidity and mortality from multiple causes ^1-3^ but is influenced by complex sets of interacting genetic loci (h^2^ >0.3-0.8) ^4,5^ that may be different between ancestries ^1,6^. Understanding the genetic basis of PLT offers important insights into the pathophysiology of PLT-mediated cardiovascular disease and disparities in health outcomes between populations ^7,8^.

Large genetic association studies for PLT and other blood cell traits have historically been restricted to European (EUR) populations ^9^. However, utilisation of non-EUR populations can reveal novel genetic associations because of differences in allele frequency, linkage disequilibrium (LD) structure and variant effects driven by environmental selection pressures and genetic drift ^10^. Trans-ethnic and ancestry-specific genome-wide association studies (GWAS) have already exploited this to reveal multiple new loci for PLT ^1,6^. Here we extend this approach by performing a GWAS in a UK collection of individuals from Bangladesh where the population prevalence of thrombocytopenia (PLT <150 ×10^9^/L) is reportedly higher than elsewhere in the Indian sub-continent ^11^, alongside a comparator population of individuals from Pakistan.

## Subjects and Methods

Analyses were performed on the July 2021 data release of the Genes and Health study (G&H) ^12^ in accordance with ethical approval from the London South-East NRES Committee of the Health Research Authority (14/LO/1240). The analysis populations comprised predominantly first-generation British Bangladeshi (BAN) and British Pakistani (PAK) adults genotyped using the Illumina Infinium Global Screening Array v3 chip.

Phenotype data were derived from linked electronic health records (EHR) which included blood cell counts measured using a Sysmex XE-2100 analyser (Sysmex, Kobe, Japan). PLT for each individual was defined as the mean of all PLT recorded in the EHRs. PLT were adjusted for sex, age, height and weight and then rank-based inverse normal transformation applied. Associations between PLT and imputed variants were calculated using BOLT-LMM v2.3.6 using the first 10 genetic principal components as covariates. Index variants were defined as those with the lowest p-value within a genome-wide significant (p <5 ×10^−8^) locus. Variants in the BAN association interval defined by index variant rs946528 underwent functional annotation using Variant Effect Predictor (VEP), mapping to epigenetic data from megakaryocytes and co-localisation with whole blood eQTLs (**Supplementary methods)**.

## Results

The characteristics of the 20,218 BAN and 9,198 PAK individuals in the analysis populations are summarised in **Table S1**. The mean PLT was lower in the BAN (mean 266.4 ×10^9^/L) than in the PAK individuals (271.5 ×10^9^/L; p = 1.16×10^−10^; **Figure S1**). SNP-based heritability for PLT was in 26.9% in BAN and 25.3% in PAK. There was high genetic correlation between the groups (r_g_ 0.94, SE 0.16).

In the BAN and PAK GWAS the genomic inflation factors were 1.096 and 1.047, respectively, indicating adequate control of population stratification. In the BAN individuals, 1588 variants assigned to 20 loci were associated with PLT at a genome-wide significance threshold of p <5 ×10^−8^ (**Figure 1, Table 1**). For each locus we next identified all variants in LD (r^2^ ≥0.5 calculated from the Bengali in Bangladesh 1000 Genomes dataset) with the index variant. The association intervals for 14 loci included at least one variant also associated with PLT in the largest to-date transethnic meta-GWAS of blood cell traits, comprising 721,201 individuals, or ancestry specific sub-analyses within this ^1^ (**Table 1**). For the remaining 6 loci, there were PLT-associated variants outside the association interval, but within 500kb of the BAN index variant, likely implicating the same locus (**Table 1**). For all 20 loci, the BAN index variants occurred at approximately the same frequency in other ancestries in the 1000 Genomes Continental genomes dataset (**Table S2**). In the PAK individuals, an identical analysis pipeline identified five significantly associated PLT loci, all of which reproduced associations found in the BAN individuals and multiple other ancestries (**Figure S2, Table S3**) ^1^.

**Figure 1:**
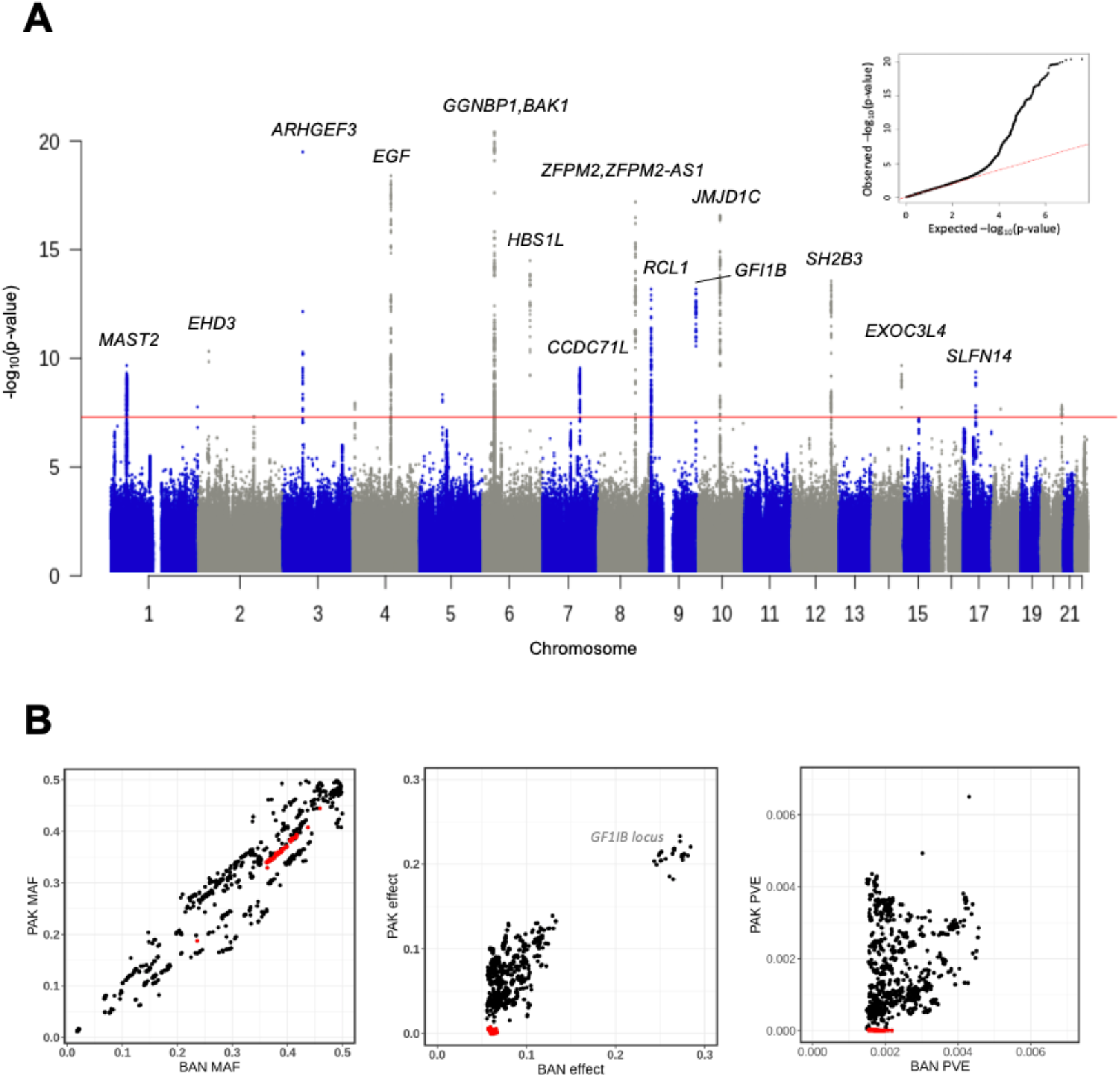
Genomic associations with PLT in 20,292 BAN individuals. **A**. Manhattan plot showing 20 loci in which the index variant has a probability of association above the genome-wide significance threshold of p <5 ×10^−8^ (red line). Selected loci are annotated with the nearest gene names identified using Variant Effect Predictor. The inset figure is the quantile-quantile plots of GWAS p-values. **B**. Minor allele frequency (MAF), effect size (beta) and phenotypic variation explained (PVE) for each of the 1596 PLT-associated variants in either BAN or PAK analyses. For the BAN locus on chromosome 1 variants in the association interval (r^2^ >0.5) of the index variant rs964528 are indicated by the red dots.

**Table 1.** Camparison of PLT-associated index variants identified in BAN individuals (n = 20,218) with index variants in previous transethnic and ancestry-specific GWAS (n = 721,201). BAN index variants were defined as those with lowest p-value within each associated locus. al positions are expressed relative to the GRCh38 genome assembly with the coded/alternate alleles on the + strand. Variants in the meta-GWAS or ancestry-specific analyses reported in Chen *et al*. ^1^ were considered indicative of the same locus as the BAN index hey were in the same LD block (r^2^ >0.5; calculated using NIH LDpop Tool with LD reference dataset from the 1000G Bengali from population (n = 86)) or within 500kb of the BAN index variants. Data are presented for the variants identified in Chen *et al*. in highest LD responding BAN variants. Trans = transethnic, EUR = European, EAS = East Asian, AFR = African, SAS = South Asian.

| BAN analysis |  |  |  |  |  |  | Transethnic meta-GWAS<sup>1</sup> |  |  |  |  |
| --- | --- | --- | --- | --- | --- | --- | --- | --- | --- | --- | --- |
| Chromosomal position | rsID | Nearest gene(s) | Coded/alternate allele | Alternate allele freq | Beta (SE) | p value | Ancestry analysis | Chromosomal position | rsID | LD between variants ($r^2$) | Distance between variants (bp) |
| chr1:46019890 | rs946528 | MAST2 | C/T | 0.58 | 0.067 (0.011) | 2.1 x10<sup>-10</sup> | EAS | chr1:45621905 | rs3014242 | 0.63 | 397,985 |
| chr1:247549001 | rs41315846 | GCSAML | T/C | 0.41 | 0.063 (0.011) | 1.7 x10<sup>-08</sup> | Trans, EUR, EAS | chr1:247556467 | rs56043070 | 0.02 | 7,466 |
| chr2:31258101 | rs592039 | EHD3 | G/A | 0.85 | 0.106 (0.016) | 4.7 x10<sup>-11</sup> | Trans, EUR | chr2:31254972 | rs655029 | 1.0 | 3,129 |
| chr2:159926221 | rs1877194 | PLA2R1 | A/G | 0.46 | -0.056 (0.010) | 4.7 x10<sup>-8</sup> | Trans, EUR, EAS | chr2:159848974 | rs12469306 | 0.17 | 77,247 |
| chr3:56815721 | rs1354034 | ARHGEF3 | T/C | 0.50 | 0.093 (0.010) | 3.2 x10<sup>-20</sup> | Trans, EUR, EAS, AFR, SAS, | chr3:56815721 | rs1354034 | Same | Same |
| chr4:6889792 | rs11734132 | KIAA0232, TBC1D14 | G/C | 0.19 | 0.079 (0.014) | 1.1 x10<sup>-8</sup> | Trans, EUR, EAS | chr4:6889728 | rs11731274 | 0.75 | 64 |
| chr4:110027510 | rs80079941 | EGF, ELOVL6 | G/C | 0.18 | -0.118 (0.013) | 4.0 x10<sup>-19</sup> | Trans, EUR, EAS | chr4:110035176 | rs76375912 | 0.97 | 7,666 |
| chr5:66710497 | rs59596869 | MAST4 | C/T | 0.13 | 0.089 (0.015) | 4.6 x10<sup>-09</sup> | Trans, EUR | chr5:66620956 | rs10940072 | 0.04 | 89,541 |
| chr6:33575632 | rs210139 | BAK1, GGNBP1 | A/C | 0.71 | 0.103 (0.011) | 3.9 x10<sup>-21</sup> | Trans, EUR, EAS, AFR | chr6:33582248 | rs210148 | 0.97 | 6,616 |
| chr6:135100038 | rs34164109 | HBS1L | C/T | 0.11 | 0.130 (0.017) | 3.2 x10<sup>-15</sup> | Trans, EUR, EAS, SAS | chr6:135101158 | rs11759553 | 1.0 | 1,120 |
| chr7:106700379 | rs342244 | CCDC71L | T/G | 0.35 | -0.069 (0.011) | 2.7 x10<sup>-10</sup> | Trans, EUR, EAS, AFR | chr7:106732176 | rs342294 | 0.86 | 31,797 |
| chr8:105570896 | rs4734879 | ZFPM2, ZFPM2-AS1 | A/G | 0.32 | -0.103 (0.012) | 6.3 x10<sup>-18</sup> | Trans, EUR, EAS | chr8:105569300 | rs6993770 | 1.0 | 1,596 |
| chr9:4788616 | rs35797651 | RCL1 | C/G | 0.35 | 0.084 (0.011) | 6.5 x10<sup>-14</sup> | Trans, EUR, EAS, SAS, HISP | chr9:4788616 | rs35797651 | Same | Same |
| chr9:132987359 | 17:355 | GFI1B | C/A | 0.02 | -0.284 (0.038) | 6.6 x10<sup>-14</sup> | Trans, EUR, SAS | chr9:132989126 | rs150813342 | <0.02 | 1,767 |
| chr10:63267383 | rs7098181 | JMJD1C | G/T | 0.48 | 0.085 (0.010) | 2.6 x10<sup>-17</sup> | Trans, EUR, EAS, AFR | chr10:63336490 | rs10822159 | 1.0 | 69,107 |
| chr12:111411711 | rs7309325 | SH2B3 | G/T | 0.41 | 0.077 (0.010) | 2.8 x10<sup>-14</sup> | EUR, EAS, AFR | chr12:111472786 | rs111818253 | 0.95 | 61,075 |
| chr14:103098397 | rs61007561 | EXOC3L4 | A/AG | 0.29 | 0.072 (0.011) | 2.1 x10<sup>-10</sup> | Trans, EUR, EAS | chr14:103100498 | rs2297066 | 1.0 | 2,101 |
| chr17:35563315 | rs55910622 | SLFN14 | G/T | 0.07 | 0.124 (0.020) | 4.2 x10<sup>-10</sup> | Trans, EUR, EAS | chr17:35558885 | rs7503168 | 0.32 | 4,430 |
| chr18:23141009 | rs11082304 | CABLES1 | G/T | 0.46 | -0.057 (0.010) | 2.1 x10<sup>-08</sup> | Trans, EUR, EAS | chr18:23141009 | rs11082304 | Same | Same |
| chr20:58999408 | rs163787 | CTSZ, NELFCD | A/G | 0.80 | 0.076 (0.013) | 1.4 x10<sup>-08</sup> | Trans, EUR, EAS | chr20:59022590 | rs46070696 | 0.02 | 23,182 |

The allele frequencies, effect sizes and phenotypic variation explained (PVE) of the PLT-associated variants identified in the BAN and PAK individuals were in most cases concordant. However, variants in the association interval at the Chromosome 1 BAN locus defined by index variant rs946528 had a disproportionately greater PVE and effect size in BAN compared with PAK individuals, whereas the allele frequencies were similar (**Figure 1B**). Accordingly, this region contained no variants that were significantly associated with PLT in PAK individuals (**Figure S2B**). The BAN rs946528 locus was also unique in that the association interval contained variants previously associated with PLT in only the East Asian (EAS) ancestry specific GWAS of Chen *et al*. whereas all the remaining 19 BAN loci reproduced findings in transethnic meta-GWAS or EUR analyses ^1^ (**Table 1**). Two other NHGRI-EBI catalogued GWAS identified significant associations with PLT for variants elsewhere in the BAN rs946528 association interval, also restricted to EAS populations (**Table S4, Figure 2**) ^13,14^. Conditional analysis of the variants in this locus with rs946528 included as a covariate revealed no secondary signals of association (p <5 ×10^−8^), indicating that rs946528 represented a single independent locus for PLT (**Figure S3**).

**Figure 2:**
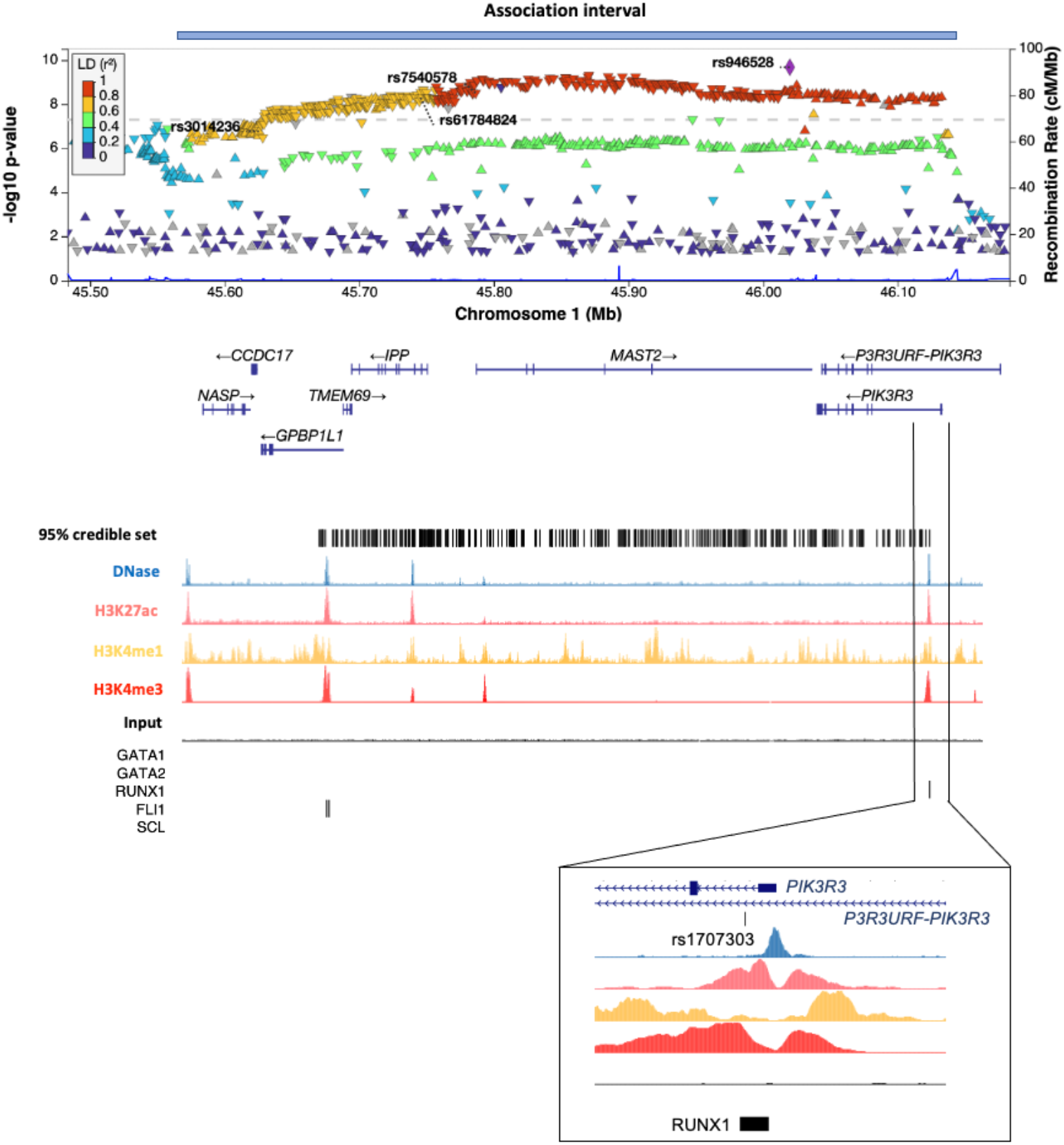
Detailed view of BAN rs946528 association interval. LocusZoom plot of associations in the 562kB association interval for the chromosome 1 index variant rs946528, defined as all significantly associated variants within r^2^ ≥0.5 calculated using LD reference data from the Bengali from Bangladesh 1000 Genomes dataset. Variants with p >0.05 are excluded to enable visualisation of the recombination peaks. Indicated variants are the index rs946528 and the three variants associated with PLT in prior EAS ancestry specific GWAS. Beneath and aligned with this are the eight UCSC Genome Browser annotated genes within the interval, the positions of 95% credible set of 343 variants and annotated regions of epigenetic activity and transcription factor binding sites from megakaryocytes, the progenitor cell for circulating platelets. The expanded view shows the relationship between the significantly associated variant rs1707303 and a putative regulatory region surrounding the first exon of the candidate gene *PIK3R3*.

The rs946528 association interval comprised a 562kB region (chr1:45575428-46137676) of low recombination containing a 95% credible set of 343 variants spanning eight UCSC Genome Browser annotated genes not previously linked to platelet phenotypes (**Figure 2, Table S5**). Within the 95% credible set, only rs1707303 in intron 1 of *PIK3R3* overlapped with a prominent area of epigenetic activity suggesting that this variant lay within a regulatory region **(Figure 2)**. *PIK3R3* also contained one of only two variants in the 95% credible set predicted to alter a canonical coding region (rs785467 predicting *PIK3R3* p.Asn283Lys). The second predicted coding region variant was rs28275469 predicting p.Lys264Arg in *IPP*, for which there were no associated variants mapped to epigenetically active areas. The rs946528 association signal colocalised with cis-eQTLs for *MAST2* and *IPP* with posterior probabilities of 45.2% and 31.4% respectively, below the threshold of 80% usually considered indicative of a shared causal variant **(Figure S5 and Table S5)**.

## Discussion

We report a large GWAS of PLT in a Bangladeshi population and an in-depth analysis of a locus on Chromosome 1 defined by the index variant rs946528. This locus was explored in more detail because the rs946528 haplotype had disproportionately larger PVE estimates in the BAN compared with PAK population for which data collection and analysis methods were identical. This finding was driven by differences in variant effect size between the BAN and PAK populations and not by MAF. The BAN rs946528 haplotype also included variants that replicated findings from other GWAS for PLT, but exclusively in EAS populations and not previously annotated ^1,13,14^. By contrast, the other PLT-associated loci in the BAN and PAK GWAS replicated prior transethnic or multiple ancestry specific GWAS, and in most cases were linked to genes already implicated in platelet biology ^1^. Replication of the rs946528 haplotype as an apparently ancestral East Asian PLT-associated locus was unsurprising given that modern BAN populations are predominantly of South Asian ancestry, but with significant East and South-East Asian admixture ^15^. This confirmatory discovery in the BAN population highlights the value of ancestry specific GWAS ^10^, in this case identifying a new PLT-associated locus that was not resolved from transethnic or large EUR population GWAS ^1,9^.

One significant challenge with the rs946528 locus is that the single independent association signal for PLT was attributable to 95% credible set of 343 variants in a haplotype block containing eight annotated genes. Considering orthogonal evidence from several sources *PIK3R3* was identified as the most likely candidate mediator of the PLT phenotype. This was primarily because the *PIK3R3* intron 1 variant rs1707303 was unique within the rs946528 haplotype in that it mapped to an area with multiple epigenetic signals indicating a *PIK3R3* regulatory region. Specifically, rs1707303, was immediately adjacent to a consensus binding site for RUNX1, a critical transcription factor necessary for differentiation and maturation of platelet forming megakaryocytes ^16^. *PIK3R3* is further supported as a candidate mediator of the PLT phenotype because it encodes phosphatidylinositol 3-kinase (PI3K) regulatory subunit [Uniprot Q92569], a regulator of PI3K in haemopoietic cells that is an essential mediator of MK maturation and platelet production ^17^.

The identification of the rs946528 haplotype as a PLT locus is directly relevant for the BAN population because cardiometabolic disease has increasing incidence in Bangladesh and particularly in Bangladeshis living in Western regions who have poorer outcomes compared to native populations ^8^. The emergence of *PIK3R3* as a candidate mediator of PLT is particularly significant as the encoded protein PI3K regulatory subunit gamma is a mediator of vascular smooth muscle proliferation and neointimal formation ^18,19^ that are critical steps in atherogenesis. This raises the possibility that the rs946528 haplotype may be a broader risk allele for cardiometabolic disease mediated by multiple cellular mechanisms, that may be tractable to therapies that target the PI3K pathway directly ^20^.

## Data availability statement

GWAS summary statistics are available for download via the journal website.

## Acknowledgements

The Genes and Health Research Team thank Social Action for Health, Centre of The Cell, members of our Community Advisory Group, and staff who have recruited and collected data from volunteers. We thank the NIHR National Biosample Centre (UK Biocentre), the Social Genetic & Developmental Psychiatry Centre (King’s College London), Wellcome Sanger Institute, and Broad Institute for sample processing, genotyping, sequencing and variant annotation. They also thank: Barts Health NHS Trust, NHS Clinical Commissioning Groups (City and Hackney, Waltham Forest, Tower Hamlets, Newham, Redbridge, Havering, Barking and Dagenham), East London NHS Foundation Trust, Bradford Teaching Hospitals NHS Foundation Trust, Public Health England (especially David Wyllie), Discovery Data Service/Endeavour Health Charitable Trust (especially David Stables), NHS Digital -for GDPR-compliant data sharing backed by individual written informed consent. Most of all we thank all of the volunteers participating in Genes & Health.

The current Genes & Health Research Team (in alphabetical order by surname) are Shaheen Akhtar, Mohammad Anwar, Elena Arciero, Omar Asgar, Samina Ashraf, Gerome Breen, Raymond Chung, Charles J Curtis, Shabana Chaudhary, Maharun Chowdhury, Grainne Colligan, Panos Deloukas, Ceri Durham, Faiza Durrani, Fabiola Eto, Sarah Finer, Ana Angel Garcia, Chris Griffiths, Joanne Harvey, Teng Heng, Qin Qin Huang, Matt Hurles, Karen A Hunt, Shapna Hussain, Kamrul Islam, Ben Jacobs, Ahsan Khan, Amara Khan, Cath Lavery, Sang Hyuck Lee, Robin Lerner, Daniel MacArthur, Daniel Malawsky, Hilary Martin, Dan Mason, Mohammed Bodrul Mazid, John McDermott, Sanam McSweeney, Shefa Miah, Sabrina Munir, Bill Newman, Elizabeth Owor, Asma Qureshi, Samiha Rahman, Nishat Safa, John Solly, Farah Tahmasebi, Richard C Trembath, Karen Tricker, Nasir Uddin, David A van Heel, Caroline Winckley, John Wright.

DV is a member of the Health Protection Research Unit in Chemical and Radiation Threats and Hazards, a partnership between Public Health England and Imperial College London which is funded by the National Institute for Health Research (NIHR).

## Author contributions

KB performed the analyses and co-wrote the manuscript, LF and DvH contributed to the analysis, DV co-designed the analyses, AM conceived the study and co-wrote the manuscript.

## Funding statement

KB is funded by a GW4-CAT Wellcome Trust Fellowship (216277/Z/19/Z). For the purpose of Open Access, the author has applied a CC BY public copyright licence to any Author Accepted Manuscript version arising from this submission.

Genes & Health is/has recently been core-funded by Wellcome (WT102627, WT210561), the Medical Research Council (UK) (M009017), Higher Education Funding Council for England Catalyst, Barts Charity (845/1796), Health Data Research UK (for London substantive site), and research delivery support from the NHS National Institute for Health Research Clinical Research Network (North Thames). Genes & Health is/has recently been funded by Alnylam Pharmaceuticals, Genomics PLC; and a Life Sciences Industry Consortium of Bristol-Myers Squibb Company, GlaxoSmithKline Research and Development Limited, Maze Therapeutics Inc, Merck Sharp & Dohme LLC, Novo Nordisk A/S, Pfizer Inc, Takeda Development Centre Americas Inc.

## Ethical approval

London South-East NRES Committee of the Health Research Authority (14/LO/1240).

## Competing interests

The authors declare no competing interests

## Notes

### Competing Interest Statement

The authors have declared no competing interest.

