## Supplementary information for "*PIK3R3* is a candidate regulator of platelet count in people of Bangladeshi ancestry"


| | |
|---|---|
| **Supplementary methods** | Page 2 |
| **Figure S1** | Page 5 |
| **Figure S2** | Page 6 |
| **Figure S3** | Page 7 |
| **Figure S4** | Page 8 |
| **Figure S5** | Page 9 |
| **Table S1** | Page 10 |
| **Table S2** | Page 11 |
| **Table S3** | Page 12 |
| **Table S4** | Page 13 |
| **Table S5** | Page 14 |
| **Supplementary references** | Page 15 |

### Supplementary methods

#### Genotyping, quality control and imputation

Genotyping, quality control and imputation were performed by the Genes and Health study. Analysis was performed on the July 2021 data release, containing 44,396 individuals genotyped on the Illumina Infinium Global Screening Array-24 v3.0 BeadChip. Quality control of genotyped data was undertaken in Ilumina GenomeStudio and plink v1.9, to remove variants with low call rate (<0.99), rare variants with minor allele frequency (MAF) <0.01, and variants that failed the Hardy–Weinberg test ($p < 1 \times 10^{-6}$). Imputation was undertaken on the TOPMed-r2Minimac4 1.5.7 Imputation Server [2].

#### Principal component analysis for genetic inference of ethnicity

Analysis was performed by Teng Heng, Wellcome Sanger Institute. Related individuals (second degree or closer) were identified using KING v2.2.4 [3], then principal component analysis (PCA) performed in 3433 unrelated reference individuals to identify distinct Pakistani and Bangladeshi clusters. The remainder of unrelated individuals were projected onto the same PC space to determine ethnicity. Ethnic outliers and individuals with discrepant questionnaire reported ethnicities were excluded from the analysis.

#### GWAS

Data preparation, GWAS and downstream analyses were performed in the Genes and Health Trusted Research Environment. Of the resulting 44,190 genotyped individuals, platelet counts (PLT) were available for 30,496 through linked electronic health records (EHRs). PLT for each individual was calculated as the mean of all recorded PLT. PLT were adjusted for age, sex, height and weight (imputed with k-Nearest Neighbour imputation where missing) using a linear regression model, outliers (>3*IQR from median) excluded, and rank-based inverse normal transformation applied to the residuals. Association statistics were calculated using BOLT-LMM v2.3.6 [4] using the first 10 PCs as covariates.

Phenotypic effect sizes were calculated as the absolute additive change in the trait mean measured in standard deviations per allele. Phenotypic variation explained (PVE) per variant was calculated as 2(MAF)*(1-MAF)*beta^2. Chromosomal positions were expressed relative to the GRCh38 genome assembly with the coded/alternate alleles on the plus strand. Variants were annotated with rsIDs using dbSNP build 155 and gene annotations using Ensembl Variant Effect Predictor (VEP) v104.3. Index variants were defined as those with the lowest p-value within a genome-wide significant ($p < 5 \times 10^{-8}$) locus.

**SNP-based heritability and genetic correlation**

SNP-based heritability and genetic correlation were estimated using LDSC v1.0.1 [5] using CSA (Central/South Asian) LD scores downloaded from the Pan-UK Biobank (https://pan.ukbb.broadinstitute.org/).

**Analysis of the rs946528 association region**

Conditional tests of association were performed using SNPtest v2.5.2 [6]. Variant-phenotype associations were conditioned upon the index variant rs946528, with repeated iterations including independently associated variants in a frequentist additive model until no associations remained ($p < 5 \times 10^{-8}$).

Statistical fine mapping was performed using FINEMAP v1.3.1 [7]. Input windows were defined as +- 500 kb from the index variant rs946528 (chr1:45,519,890-46,519,890). The number of conditionally independent signals in the window was used as prior knowledge for the maximum number of causative variants to be searched (–n-causal-snps option). The LD structure was computed from the same samples included in the GWAS analysis. 95% credible sets were defined as minimal sets of variants jointly covering at least 95% of the posterior probability of including the true causative signal.

Predicted regulatory regions were interrogated in the UCSC Genome Browser (http://genome.ucsc.edu/), using DNase I hypersensitivity, H3K27ac (active gene transcription), H3K4me1 (enhancer) and H3K4me3 (promoter) data tracks from CD34-negative, CD41-positive, CD42-positive megakaryocytes provided by the BLUEPRINT Epigenomics Project [8]. ChIP-seq identified binding sites for haematopoietic transcription factors GATA1, GATA2, RUNX1, FLI1, and SCL in megakaryocytes were also integrated [9].

Colocalisation of the GWAS signal with eQTL datasets was performed using Approximate Bayes Factor analysis in the R package coloc (https://cran.r-project.org/web/packages/coloc/). Whole blood summary cis-eQTL data was downloaded from the eQTLGen Consortium https://www.eqtlgen.org/ [10] and posterior probability for both traits (PLT versus eQTL) sharing a single causal variant ($H_4$) calculated for *NASP, CCDC17, GPBP1L1, TMEM69, IPP, MAST2* and *PIK3R3*.

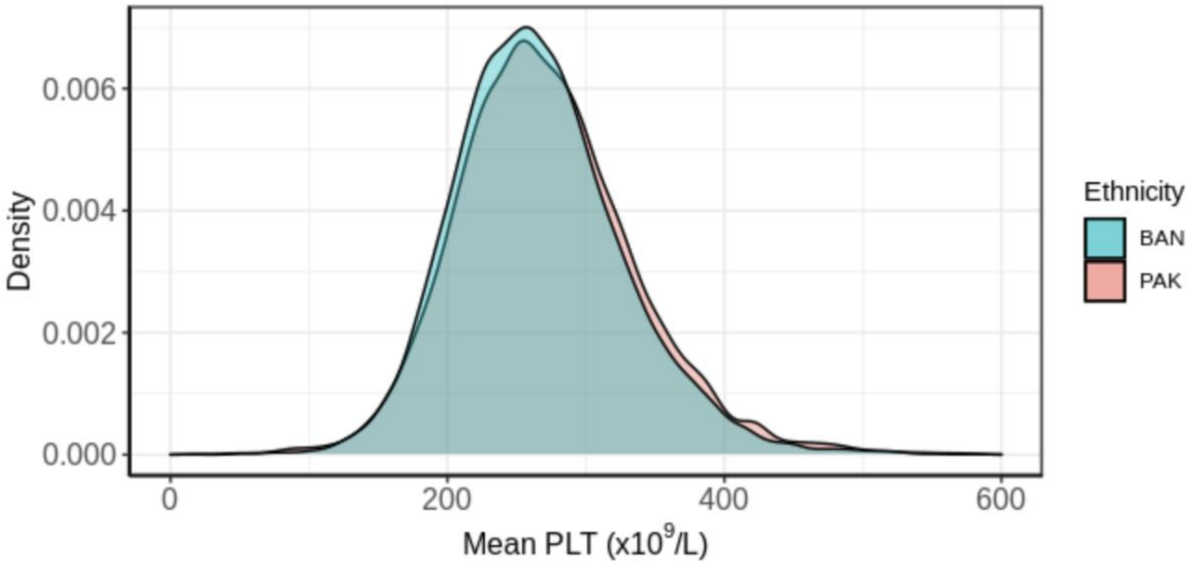


**Figure S1: PLT in the BAN and PAK individuals**

For each individual in the analysis groups, the data represent the mean of all PLT documented in electronic health records within the Genes and Health dataset.

**A**

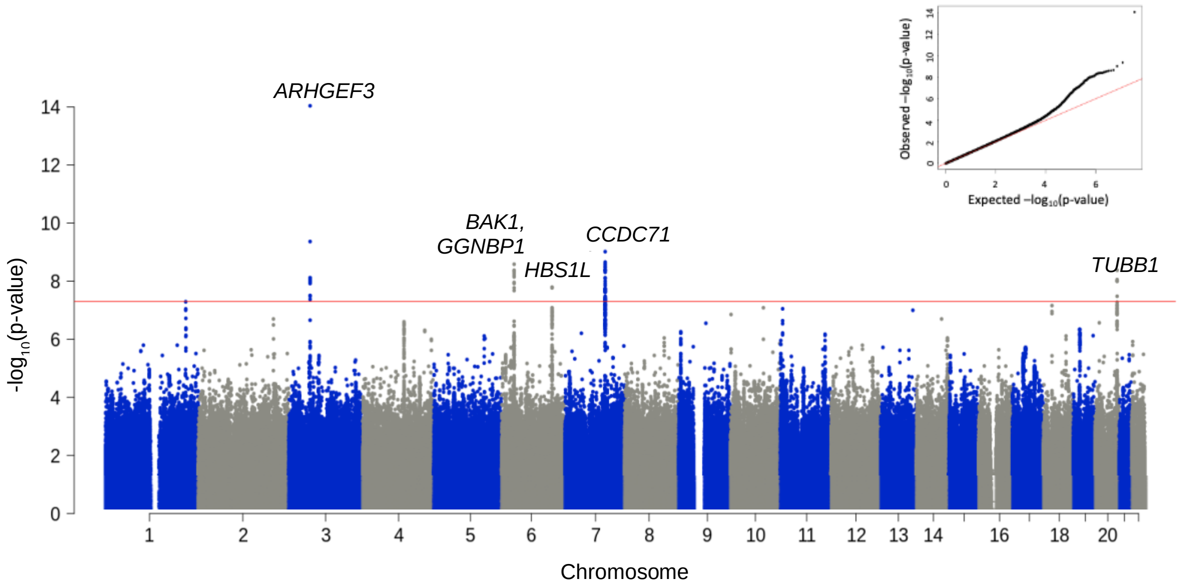


**B**

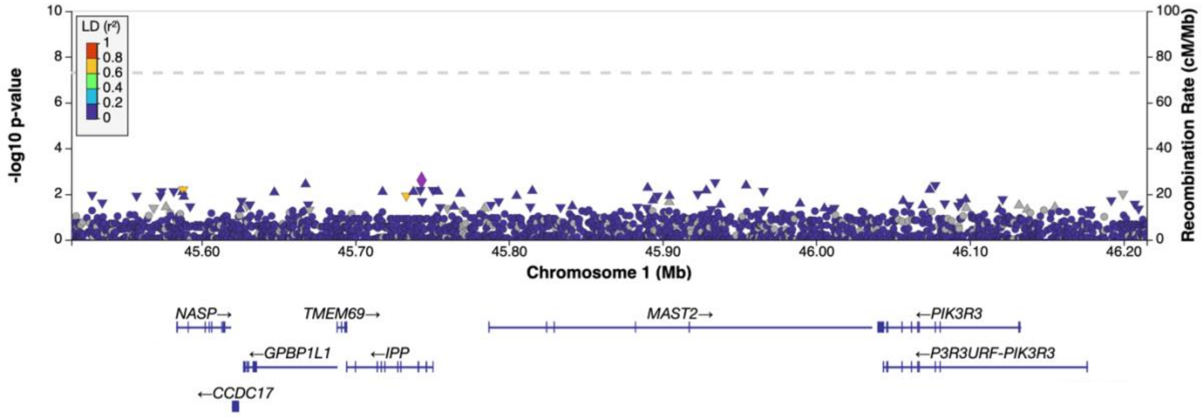


**Figure S2: Associations between variants and PLT in 9,198 PAK individuals**

**A**. Manhattan plot of genomic associations with PLT showing five significantly associated loci annotated with the names of the nearest genes identified using VEP. The red line indicates the genome-wide significance threshold of p <5 x10$^{-8}$. The inset figure is the quantile-quantile plot of GWAS p-values. **B.** Regional plot of the r946528 association interval in PAK individuals. Variants in this interval were significantly associated with PLT in BAN individuals, but none reached genome wide significance (dashed line) in PAK individuals.

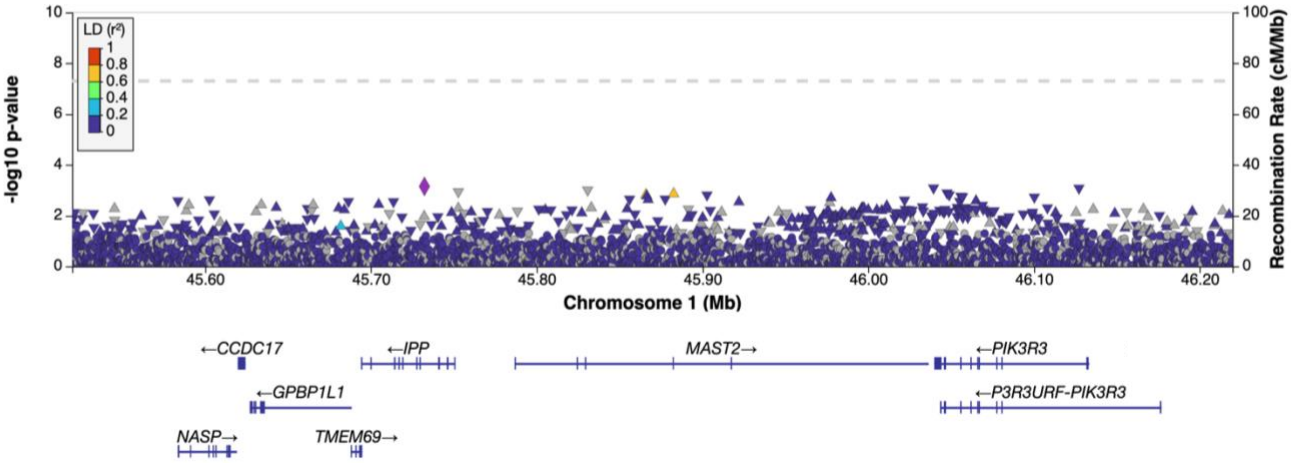


**Figure S3: Conditional analysis of genomic associations with PLT in the rs946528 association interval in 20,292 BAN individuals.** There were no secondary signals of association for variants in this region when the index variant rs946528 was included as a covariate in the analysis. The labelled variant (purple diamond) has the lowest p-value in the region.

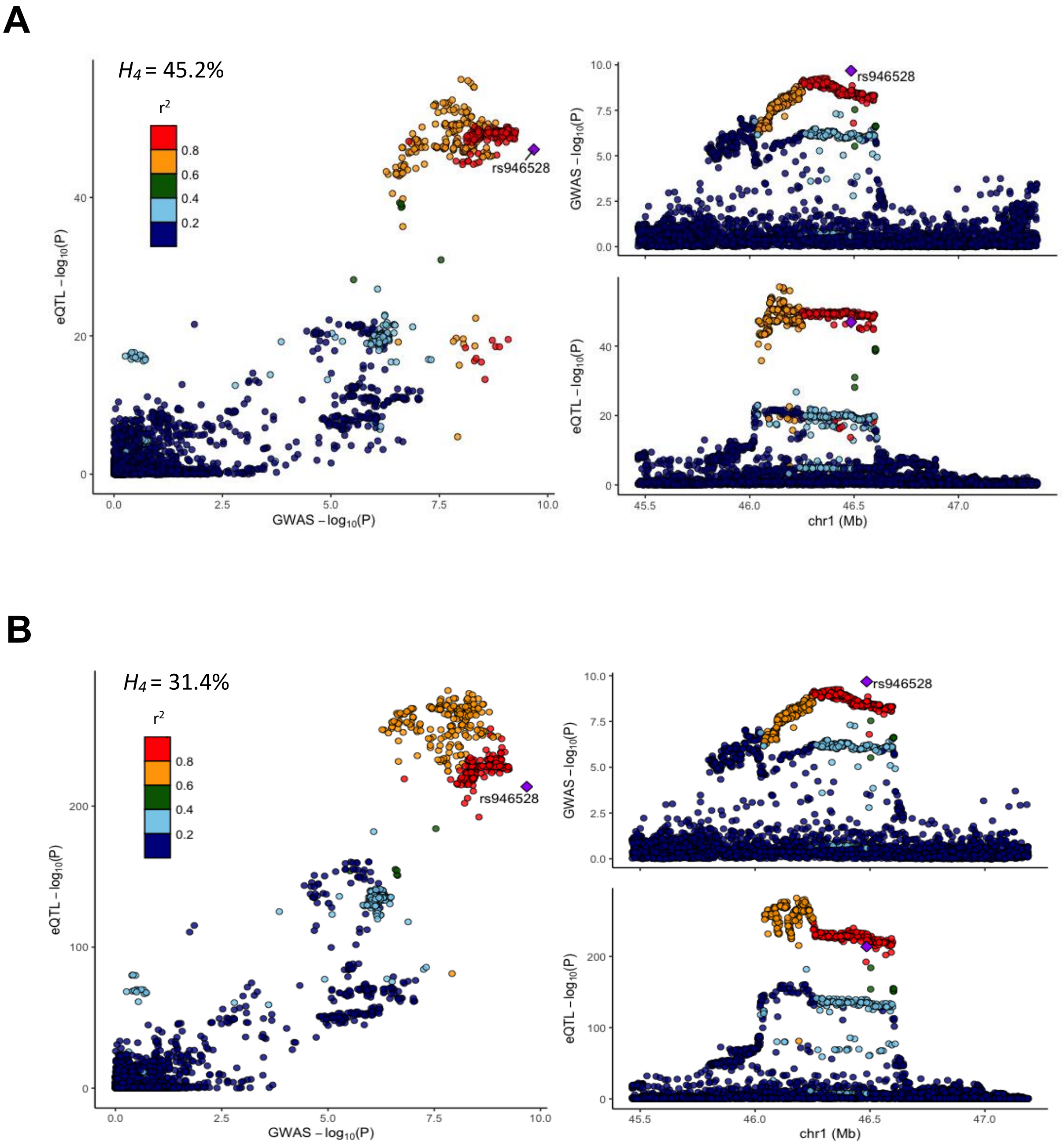


**Figure S5: Co-localisation of variants in the r946528 association interval with whole blood eQTLs for proximal genes.**

Data are presented for (**A**) *MAST2* and (**B**) *IPP* eQTLs. Posterior probabilities of a single causal variant ($H_4$) were calculated using *coloc.abf()* function of R package *coloc* and plotted using *locuscomparer*. The main plots show correlation between p-values for GWAS and eQTL variants, with the inset plots showing regional views for both datasets.

**Table S1: Clinical Characteristics of the study populations.** Data are reported as mean (standard deviation).

| | **BAN (n = 20,218)** | **PAK (n = 9,198)** |
|---|---|---|
| Gender (male) | 43.4% | 41.4% |
| Age at recruitment (years) | 41.0 (12.9) | 44.6 (15.3) |
| Height (cm) | 160 (8.5) | 164 (9.1) |
| Weight (Kg) | 67.2 (12.3) | 74.4 (14.9) |

**Table S2: Allele frequencies of PLT-associated variants from BAN analysis and in 1000 Genomes Continental Populations.** AFR = African, AMR = Ad Mixed American, EAS = East Asian, EUR = European, SAS = South Asian.

| Chromosomal position (GRCh38) | rsID | Nearest gene(s) | Coded/alternate allele | | | Superpopulation alternate allele frequency | | | | |
|---|---|---|---|---|---|---|---|---|---|---|
| | | | | BAN | PAK | AFR | AMR | EAS | EUR | SAS |
| chr1:46019890 | rs946528 | MAST2 | C/T | 0.58 | 0.61 | 0.53 | 0.61 | 0.71 | 0.72 | 0.56 |
| chr1:247549001 | rs41315846 | GCSAML | T/C | 0.41 | 0.40 | 0.70 | 0.33 | 0.45 | 0.50 | 0.41 |
| chr2:31258101 | rs592039 | EHD3 | G/A | 0.85 | 0.81 | 0.89 | 0.84 | 0.94 | 0.69 | 0.89 |
| chr2:159926221 | rs1877194 | PLA2R1 | A/G | 0.46 | 0.48 | 0.46 | 0.77 | 0.61 | 0.77 | 0.43 |
| chr3:56815721 | rs1354034 | ARHGEF3 | T/C | 0.50 | 0.53 | 0.20 | 0.44 | 0.57 | 0.60 | 0.48 |
| chr4:6889792 | rs11734132 | KIAA0232, TBC1D14 | G/C | 0.19 | 0.14 | 0.11 | 0.17 | 0.53 | 0.17 | 0.16 |
| chr4:110027510 | rs80079941 | EGF, ELOVL6 | G/C | 0.18 | 0.13 | 0.05 | 0.01 | 0.24 | 0.00 | 0.19 |
| chr5:66710497 | rs59596869 | MAST4 | C/T | 0.13 | 0.06 | 0.16 | 0.01 | 0.04 | 0.01 | 0.13 |
| chr6:33575632 | rs210139 | BAK1, GGNBP1 | A/C | 0.71 | 0.70 | 0.52 | 0.53 | 0.73 | 0.42 | 0.74 |
| chr6:135100038 | rs34164109 | HBS1L | C/T | 0.11 | 0.11 | 0.14 | 0.16 | 0.24 | 0.26 | 0.11 |
| chr7:106700379 | rs342244 | CCDC71L | T/G | 0.35 | 0.34 | 0.24 | 0.34 | 0.25 | 0.42 | 0.40 |
| chr8:105570896 | rs4734879 | ZFPM2, ZFPM2-AS1 | A/G | 0.32 | 0.33 | 0.42 | 0.31 | 0.40 | 0.29 | 0.36 |
| chr9:4788616 | rs35797651 | RCL1 | C/G | 0.35 | 0.22 | 0.04 | 0.22 | 0.63 | 0.23 | 0.31 |
| chr9:132987359 | rs149810016 | GFI1B | C/A | 0.02 | 0.01 | 0.00 | 0.00 | 0.00 | 0.00 | 0.04 |
| chr10:63267383 | rs7098181 | JMJD1C | G/T | 0.48 | 0.48 | 0.28 | 0.30 | 0.33 | 0.43 | 0.50 |
| chr12:111411711 | rs7309325 | SH2B3 | G/T | 0.41 | 0.35 | 0.77 | 0.32 | 0.89 | 0.20 | 0.34 |
| chr14:103098397 | rs61007561 | EXOC3L4 | A/AG | 0.29 | 0.24 | 0.20 | 0.21 | 0.26 | 0.24 | 0.31 |
| chr17:35563315 | rs55910622 | SLFN14 | G/T | 0.07 | 0.08 | 0.08 | 0.04 | 0.01 | 0.04 | 0.07 |
| chr18:23141009 | rs11082304 | CABLES1 | G/T | 0.46 | 0.49 | 0.21 | 0.36 | 0.53 | 0.51 | 0.46 |
| chr20:58999408 | rs163787 | CTSZ, NELFCD | A/G | 0.80 | 0.80 | 0.88 | 0.78 | 0.86 | 0.82 | 0.84 |

**Table S3: Comparison of PLT-associated index variants identified in PAK individuals (n = 9198) with index variants in a prior transethnic meta-GWAS (n = 721,201) [1].** PAK index variants were defined as those with lowest p-value within each associated locus. Chromosomal positions are expressed relative to the GRCh38 genome assembly with the coded/alternate alleles on the + strand. Variants in the transethnic meta-GWAS were considered indicative of the same locus as the PAK index variant if they were in the same LD block ($r^2$ >0.5 with LD reference dataset from the 1000G Punjabi from Lahore, Pakistan population (n = 96)) or within 500kb of the PAK index variant. Data are presented for the associated variant identified in Chen *et al.* in highest LD with the corresponding PAK variant. Trans = transethnic, EUR = European, EAS = East Asian, AFR = African, SAS = South Asian.

| PAK analysis | | | | | | | Transethnic meta-GWAS [1] | | | | |
|---|---|---|---|---|---|---|---|---|---|---|---|
| **Chromosomal position (GRCh39)** | **rsID** | **Nearest gene(s)** | **Coded/ alternate allele** | **Alternate allele frequency** | **Beta (SE)** | **p value** | **Ethnicity** | **Chromosomal position** | **rsID** | **LD between variants ($r^2$)** | **Distance between variants (bp)** |
| chr3:56815721 | rs1354034 | *ARHGEF3* | T/C | 0.53 | 0.114 (0.015) | $9.1 \times 10^{-15}$ | Trans, EUR, EAS, AFR, SAS | chr3:56815721 | rs1354034 | Same | Same |
| chr6:33596519 | rs12206050 | *BAK1, GGNBP1* | A/T | 0.35 | 0.091 (0.015) | $2.6 \times 10^{-09}$ | Trans, EUR, EAS, AFR | chr6:33578721 | rs5745582 | 0.98 | 17,798 |
| chr6:135110180 | rs6920211 | *HBS1L* | T/C | 0.18 | 0.130 (0.023) | $1.6 \times 10^{-8}$ | Trans, EUR, EAS, SAS | chr6:135101158 | rs11759553 | 0.95 | 9,022 |
| chr7:106721764 | rs342284 | *CCDC71L* | T/C | 0.31 | -0.101 (0.016) | $9.6 \times 10^{-10}$ | Trans, EUR, EAS, AFR | chr7:106731776 | rs342294 | 0.89 | 10,012 |
| chr20:59019629 | rs34524896 | *TUBB1* | C/T | 0.13 | -0.130 (0.022) | $4.2 \times 10^{-9}$ | Trans, EUR, EAS | chr20:59022590 | rs6070696 | 0.03 | 2,961 |

**Table S4: Variants in the BAN rs946528 association interval identified as PLT-associated loci in previous GWAS.** LD was calculated using NIH LDpop Tool with LD reference dataset from the 1000G Bengali from Bangladesh population (n = 86)).

| Chromosomal position (GRCh38) | rsID | Ancestry | VEP annotation | LD with rs946528 ($r^2$) | Distance to rs946528 (bp) | Reference |
|---|---|---|---|---|---|---|
| chr1:46019890 | rs946528 | BAN | *MAST2* intronic | - | - | Current study |
| chr1:45745675 | rs61784824 | EAS: Japan | *MAST2* promoter flanking | 0.87 | 274,215 | 11 |
| chr1:45621905 | rs3014242 | EAS: Japan | *IPP* intronic | 0.66 | 397,985 | 12 |
| chr1:45786472 | rs7540578 | EAS: Japan, UK, China | *CCDC17* missense | 0.63 | 233,418 | 1 |

**Table S5: Genes present in the rs946528 association interval (chr1:45575428-46137676).** The association interval is defined as the interval containing all variants in LD ($r^2$ >0.5) with index SNP rs946528. Gene names and functional annotations are derived from NCBI Gene (www.ncbi.nlm.nih.gov/gene/). Expression data for CD34-negative, CD41-positive, CD42-positive megakaryocytes from the BLUEPRINT Epigenome Project (www.blueprint-epigenome.eu/). FPKM = fragments per kilobase of transcript per million mapped reads. The posterior probability of colocalisation between variants in the BAN PLT rs946528 association interval and whole blood eQTLs for each gene in the interval are calculated using eQTLGen data (https://www.eqtlgen.org/).

| **Gene** | **Encoded protein** | **Biological function** | **Gene expression in MKs (FPKM)** | **Posterior probability of shared causal variant ($H_4$)** |
|---|---|---|---|---|
| *NASP* | Nuclear autoantigenic sperm protein | H1 histone binding protein required for nuclear histone transport that I necessary for DNA replication and cell cycle progression. | 77.645 | 0.06% |
| *CCDC17* | Coiled-coil domain containing protein 17 | Protein of unknown function predominantly expressed in lung. | 0.525 | 10.1% |
| *GPBP1L1* | GC-rich promoter binding protein 1 like 1 | Putative DNA/RNA binding protein and predicted regulator of transcription. | 80.03 | 0.46% |
| *TMEM69* | Transmembrane Protein 69 | Predicted membrane protein of unknown function. | 32.805 | 0.009% |
| *IPP* | Intracisternal A particle-promoted polypeptide | Kelch protein family member with predicted actin binding domains. | 4.95 | 31.4% |
| *MAST2* | Microtubule associated serine/threonine Kinase 2 | Microtubule associated serine/threonine kinase. | 1.54 | 45.2% |
| *PIK3R3* | Phosphatidylinositol 3-kinase regulatory subunit gamma | Regulatory component of the phosphatidylinositol 3-kinase complex that phosphorylates phosphatidylinositol and similar compounds in multiple cellular processes. | 1.28 | 0.61% |
| *P3R3URF-PIK3R3* | P3R3URF-PIK3R3 readthrough | Naturally occurring readthrough transcription between neighbouring genes LOC110117498 and *PIK3R3*. | Unknown | N/A |
